# Thymically imprinted differential responsivity of naïve CD4 T cells to IL2 results in a heterogeneity of both Treg induction and subsequent effector functioning

**DOI:** 10.1101/2023.10.08.561378

**Authors:** Nathan D. Pennock, Yamin Qian, Kazumi Ishihara, Yamami Nakamura, Eric Cross, Shimon Sakaguchi, Jason T. White

## Abstract

Thymic selection predisposes naive T cells to particular outcomes when challenged later with cognate antigen, whether the antigen is self or foreign. This suggests that there is an inherent heterogeneity of functioning among T cells within the naive population (both CD4 and CD8s), and that each T cell, as part of its thymic development, is given a certain ‘programming’ which will affect its eventual fate decisions. In this project, we looked at the primary effects of this thymic imprinting on the conversion of naïve CD4 T cells into Tregs. Further, using an induced-Treg-reporter system, we exam the impact of thymic imprinted heterogeneity on effector functionality and identity stability. We report that naïve T cell differential responsivity to cytokines leads to the observed difference in Treg induction, and that the Tregs induced from T cells of different self-affinities maintain a heterogeneity of effector function and identity.

## Introduction

A T cell’s affinity for self-peptide/MHC during thymic maturation determines a T cell’s CD4 and CD8 status, natural Treg identity, and the decision to live, die, or revise their T cell receptor through the processes of positive and negative selection (Goldrath and Bevan, 1999; Hogquist et al., 1994; Singer et al., 2008). The outcome of these processes is the generation of identifiable, thymically matured single positive T cells that exit into the periphery, and constitute the systemic naïve T cell repertoire. By virtue of passing through thymic selection, these T cells express a revised, but still broad, range of self-affinity, deemed to be both useful and safe to the host.

Historically speaking, upon reaching this stage, one naïve cell was generally considered to be equivalent to any other, with the chief exception being a cell’s antigen specificity and associated affinity for antigen. Consistent with this, phenotypically, single positive CD4 and CD8 T cells in this thymically mature/peripherally naïve state display tremendous plasticity for expressing transcription factors that determine a broad range of Th and Tc states (Th/c 1,2,17) based upon the activation conditions of the peripheral milieu when their antigen is present (Pennock et al., 2013). Results from studies into homeostasis in lymphopenic settings began to challenge this assumption, however, when it was determined that cells of a higher thymic self-affinity were better able to proliferate in the cytokine-rich environment of lymphodepleted animals (Kieper et al., 2004). The concept that there could be a heterogeneity of function within the naïve T cell pool stratified along self-affinity has steadily been applied to various aspects of CD4 and CD8 T cell functioning, and it has been determined that there are indeed a number of differences, both phenotypic and functional (Fulton et al., 2015; Persaud et al., 2014; White et al., 2016).

Perhaps the earliest and most consistent phenotypic difference, long known to tightly correlate with the thymic affinity of TCR-pMHC interactions, is the expression of the plasma membrane protein CD5. CD5 is a type I transmembrane glycoprotein composed of an extracellular region possessing three scavenger receptor cysteine rich domains (SRCR) and an intracellular tail possessing tyrosine and serine phosphorylation targets that serve as intracellular signaling scaffold. CD5 surface expression is experimentally useful both for the fact that it directly correlates with the amount of signaling a T cell receives during thymic selection and this level appears to be fixed for the duration of the cell’s naïve state (Azzam et al., 1998). Functionally, the absence of CD5 in thymic T cells prior to the DP state results in enhanced TCR signaling, shifting the affinity windows of positive and negative selection downward (Tarakhovsky et al., 1995). In the periphery, CD5 expression can be altered by activation and the function of CD5 is unclear, as a variety of treatment conditions result in varied states of intracellular tail phosphorylation and recruitment of a diversity of scaffold binding partners (Burgueno-Bucio et al., 2019). Due to limited interrogation tools and the tight correlation between CD5 and self-affinity, in previous reports differences in cell character due to varying self-affinities were often ascribed by assumption rather than molecular interrogation, to the functionalities of the CD5 molecule itself. In this report, we leverage modern CRISPR-CAS9 approaches to diminish CD5 expression specifically in the states of T cell interest to clearly define the role of CD5 in naïve T cell polarization.

Fitting with the tight correlation between CD5 and thymic TCR-pMHC affinity, the highest expressers of CD5 under physiological conditions are CD4+ T regulatory cells that develop in the thymus (natural or thymic Tregs [nTregs/tTregs]). nTregs have long been known to arise as a consequence of high thymic self-affinity interactions (Jordan et al., 2001; Kieback et al., 2016). As such, it is perhaps not surprising that studies into naive CD4+ T cells (Tconv) under regulatory polarization conditions have generally agreed that there is a propensity of cells with a higher self-affinity (and therefore higher CD5) to become induced or peripheral Tregs (iTreg/pTreg) (Henderson et al., 2015; Martin et al., 2013). This finding is by no means universal, nor has the reason or mechanism of this association been definitively explored. Given the known dramatic stromal determinants of cell fate in the thymus, and the radical distinction between the architecture and environments of the thymus and the periphery, it is unclear how trends in nTreg differentiation in the thymus would readily translate into similar outcomes in the periphery. Furthermore, unconfounded consideration has not been given to the to the function of CD5 itself in influencing iTreg development. In an attempt to unify the field, and to address these outstanding questions, we undertook a comprehensive study into CD5 and naïve CD4 T cells fate under conditions of Treg polarization.

Finally, previous studies into the correlation of self-affinity and T cell functioning have either focused only on the naïve phase or have studied secondary effectors as a mixed affinity population. Here, using a Foxp3 reporter system, we were able to isolate iTregs sourced from Tconv of varying self-affinities and compare their functioning as effectors on a per-cell basis. In the following studies, we determined not only that Tconv differential responsivity to cytokines leads to the observed difference in iTreg induction, but also that iTregs from Tconv of different self-affinities maintain a heterogeneity of function once they reach the effector phase in vitro and in vivo.

## Results

### CD5 correlates with Treg induction in vitro and in vivo

In our studies, we leveraged the well-established relationship between thymic self-antigen affinity and surface CD5 in order to investigate the role of thymic self-antigen affinity in programming intrinsic peripheral T cell heterogeneity. It has been demonstrated previously that not only does CD5 surface expression directly correlate with affinity for self-antigen experience in the thymus, but in addition this set level of CD5 expression seems to remain largely constant while the T cell remains in a naïve state (Azzam et al., 1998). Further, to improve upon similar efforts undertaken in the past we have guarded against unintentional skewing of results that might occur through inclusion of phenotypically naïve cells which have nascently begun to express FoxP3, through employment of sorts using transgenic mice expressing a Foxp3 reporter (either eGFP [FDG] or human CD2 [hCD2]) (Kim et al., 2007; Komatsu et al., 2009). For our initial interrogation into the relationship between self-affinity and Treg formation, we took naïve CD4 splenocytes (Lineage- /CD4+, CD44lo, CD25-Fig 1a) and sorted the 15% lowest and highest expressers of CD5 (low and high self-affinity, respectively) (Fig 1a). After three days of in vitro differentiation conditions with CD3 + CD28 beads, TGFb, and IL-2, we observed that CD5hi cells were significantly more likely to differentiate into induced Tregs (iTregs) as determined by Foxp3 protein expression (Fig 1b). To determine when the association of CD5 level with Treg inducibility is established, we repeated the induction experiment, starting with mature single-positive CD4 thymocytes (Dump-/CD4+, CD44lo, MHCI lo, CD25- Figure 1c); a population which has completed thymic selection, and is ready to egress to the periphery (Hogquist et al., 2015). We observed the same pattern in the thymocytes as was observed in naïve splenocytes, with high self-affinity cells being more predisposed to iTreg formation, thereby establishing that the predisposition of Tconv with high self-affinity to become iTreg is present at least as soon as the T cell achieves the mature, single positive thymocyte state.

**Figure 1.**
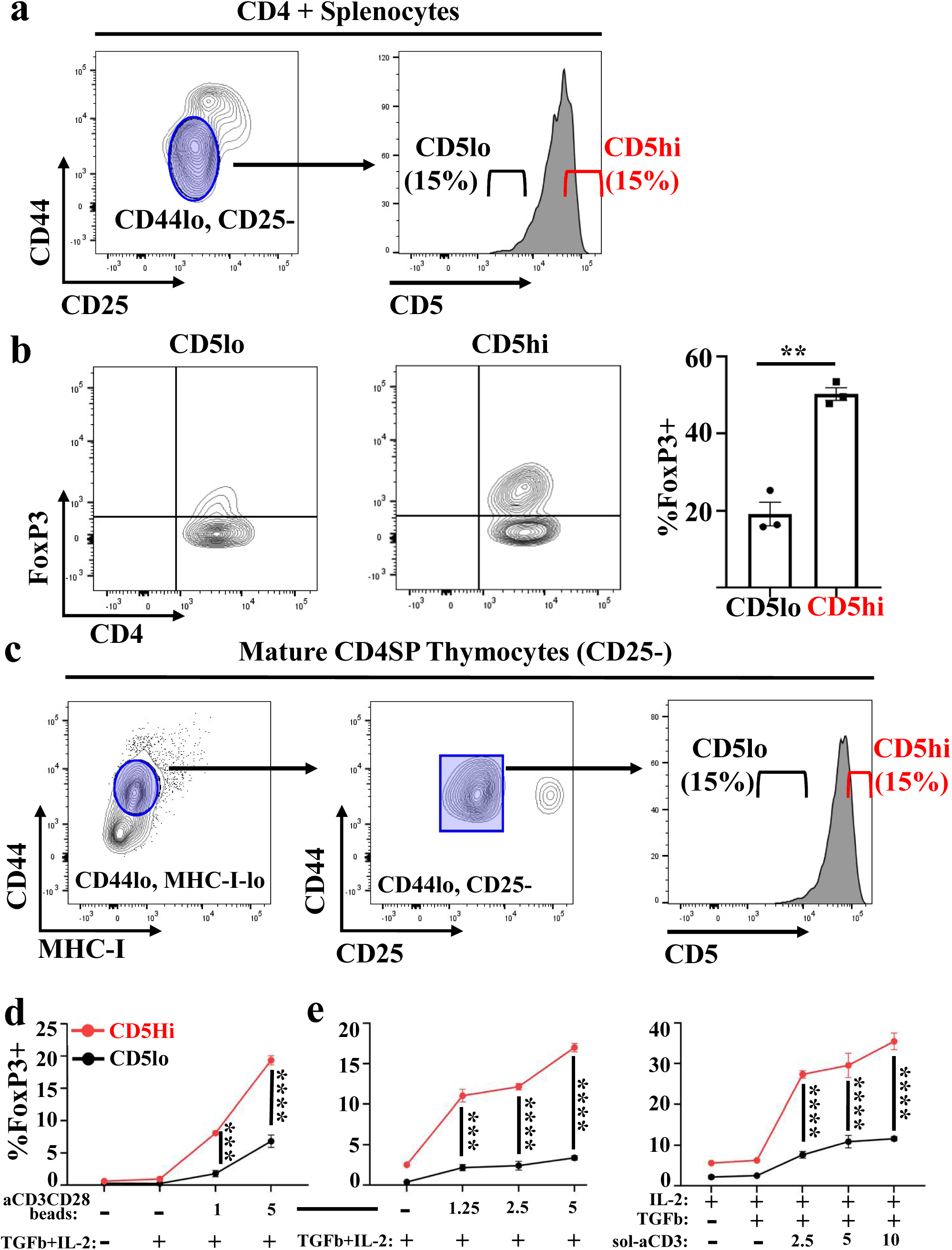
CD5 correlates with Treg induction in vitro. a. Gating strategy for obtaining CD5lo and CD5hi naïve Tconv from CD4+ splenocytes (CD4+, CD8-, B220-, Foxp3-eGFP/Foxp3-hCD2-). b. Percentage of iTreg obtained from in vitro induction experiments on CD5-disparate naïve Tconv splenocytes: aCD3/CD28 beads (5ug/mL) with IL2 (50U/mL) and TGFb (2.5ng/mL) for 3 days (n=3). Data representative of multiple experiments. c. Gating strategy for obtaining CD5lo and CD5hi mature single-positive thymocytes from CD4+ thymocytes (CD4+, CD8-, B220-, Foxp3-eGFP/Foxp3-hCD2-). d. Percentage of iTreg obtained from in vitro induction experiments on CD5-disparate mature single-positive thymocytes: varying concentrations of aCD3/CD28 beads with IL-2 (50U/mL) and TGFb (2.5ng/mL) for 3 days (n=3). Data representative of multiple experiments. e. Further experiments in the in vitro induction of CD5-disparate naïve Tconv splenocytes: varying concentrations of aCD3/CD28 or immobilized soluble aCD3 (ug/mL) with IL-2 (50U/mL) and TGFb (2.5ng/mL) for 3 days (n=3). Data representative of multiple experiments.

Importantly this demonstrates that proclivity to iTreg differentiations is not a trait acquired in the periphery (Fig 1d). Returning to the splenocytes, we next altered concentrations of reagents in the induction cocktail to determine the robustness of the predisposition of CD5hi cells to iTreg formation. We found that the predisposition held true across varying concentrations of TCR stimulation, regulatory cytokines (IL-2, left, TGFb, supplemental figure 1a), and whether or not the T cell received signaling through CD28 (aCD3/CD28 bead stimulation or aCD3 antibody alone) (Fig 1e, right). In addition to the clear superiority of CD5hi cells to generate Foxp3+ iTregs, we also noted that only TCR stimulation significantly increased Foxp3+ inducibility of CD5hi cells, and it did so in a dose dependent manner.

As our observations were robustly upheld through a variety of in vitro cell culture differentiation conditions, we next sought to determine whether thymic predetermination of cellular differentiation was truly a biologically relevant phenomenon. To do this we combined our CD5 sorting approach with an adoptive transfer model in vivo. Due to the phenomenon of CD5hi cells being more auto-reactive in lymphopenic hosts (Kieper et al., 2004), we endeavored to further improve the system by using a lymphoreplete, antigen specific model of oral tolerance. In this model, CD5lo and CD5hi splenic CD4 Tconv from OTII/RAGKO mice were isolated and adoptively transferred into congenically disparate, lymphoreplete, unmanipulated B6 mice. It is important to note that, even in a fixed, single clonotypic TCR transgenic system (such as that employed here with the OTII/RAGKO mice), Tconv show a narrower but nonetheless observable distribution of CD5 values (Fig 2a). Further, when sorted into CD5hi and CD5lo populations and differentiated in vitro (as described above), we observe a similar proclivity for CD5hi towards iTreg differentiation. (Fig 2b).

**Figure 2.**
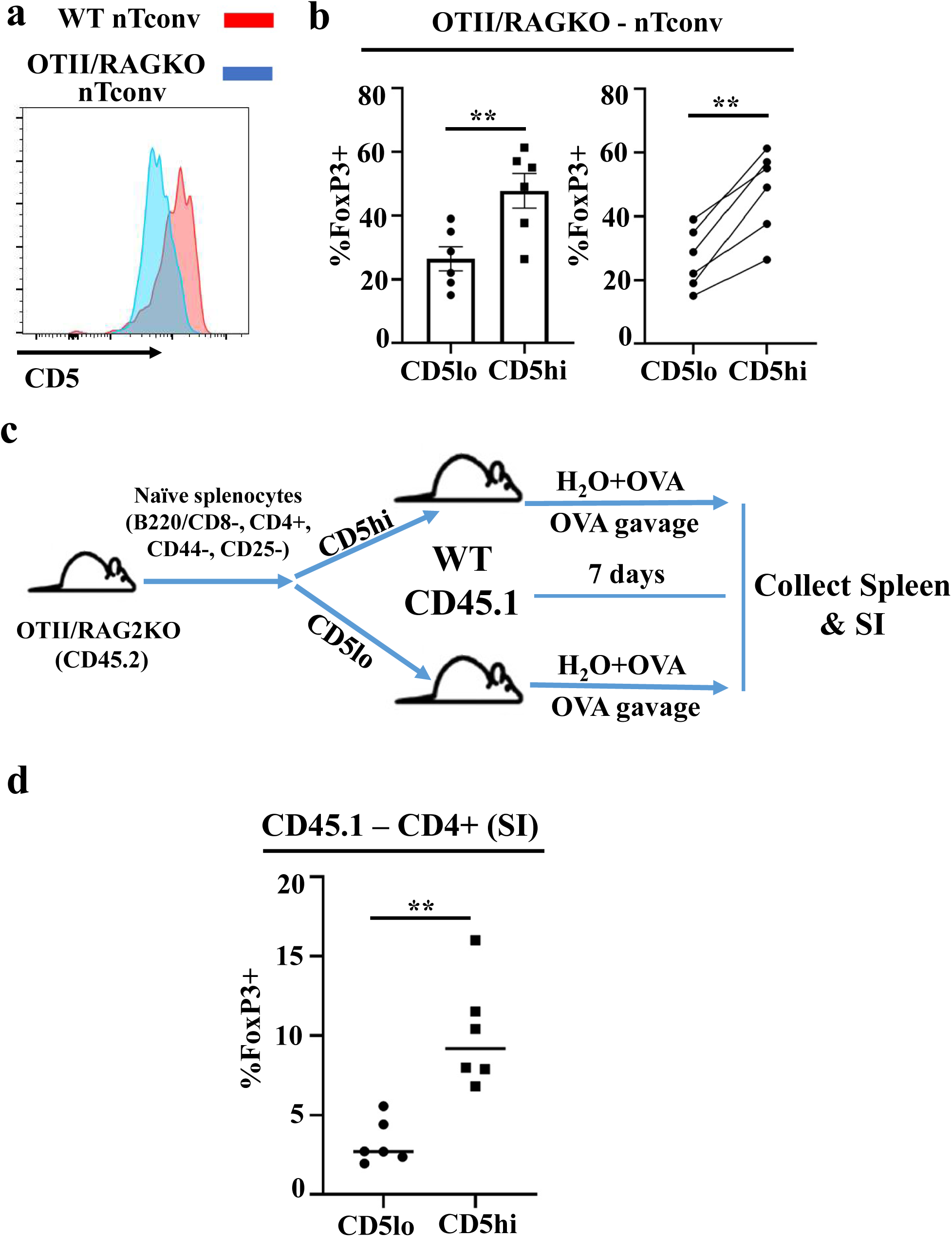
CD5 correlates with Treg induction in vivo. a. Representative distribution of CD5 on naïve Tconv splenocytes from WT and OTII/RAGKO mice. b. Percentage of iTreg obtained from in vitro induction experiments on CD5-disparate naïve Tconv splenocytes from OTII/RAGKO mice: immobilized soluble aCD3 (10ug/mL) with IL2 (50U/mL) and TGFb (2.5ng/mL) for 3 days (n=6). Same data shown in bar graph and line graph, for illustrative purposes. Data representative of multiple experiments. c. CD5lo and CD5hi Tconv splenocytes flow sorted from OTII/RAGKO mice were sorted and adoptively transferred into congenic WT mice (day 0). Recipient mice were provided water containing OVA protein (10mg OVA/mL, ad lib), and received gavage of OVA protein on days 1, 3, and 5 (20mg OVA in 200uL PBS). Mice sacrificed and tissues analyzed on day 7. d. Percentage of transferred OTII/RAGKO Tconv cells from the small intestine converting to a Foxp3+ phenotype. Data combined from two experiments (n=3/experiment).

In our system, wildtype B6 mice receiving OTII/RAGKO CD5lo/hi Tconv transfers were subsequently provided with water containing OVA protein and fed OVA by oral gavage for one week. Mice were euthanized after a week, and spleen and small intestines were collected, dissociated, and analyzed by flow cytometry to determine the relative fate of the transferred cells (Fig 2c). While there was no measurable difference in Tregs between the groups in the spleen (data not shown), in the small intestine a significantly higher proportion of the CD5hi cells were Foxp3+, establishing that the in vitro findings hold true in a lymphoreplete, antigen-specific in vivo system (Fig 2d).

### The CD5 molecule is not responsible for differences in iTreg induction

To determine the extent of the correlation between self-affinity and iTreg induction with greater resolution, we induced iTregs from splenic naïve Tconv populations flow sorted across a broader range of CD5 expression. We observed significantly tight linear correlations between CD5 and Foxp3 expression under conditions of both TCR-only stimulation and TCR/CD28 stimulation (Fig 3a). These results agreed with our prior experiments of discrete CD5 (hi/lo) assignments across cell types (thymic/splenic) and their associated differential CD5 distributions under various in vitro polarization conditions (supplemental 1b-d). The association was even maintained across TCR repertoires, which we tested by examining the induction of Tconv from our WT and OTII/RAGKO mice (Fig 3b).

**Figure 3.**
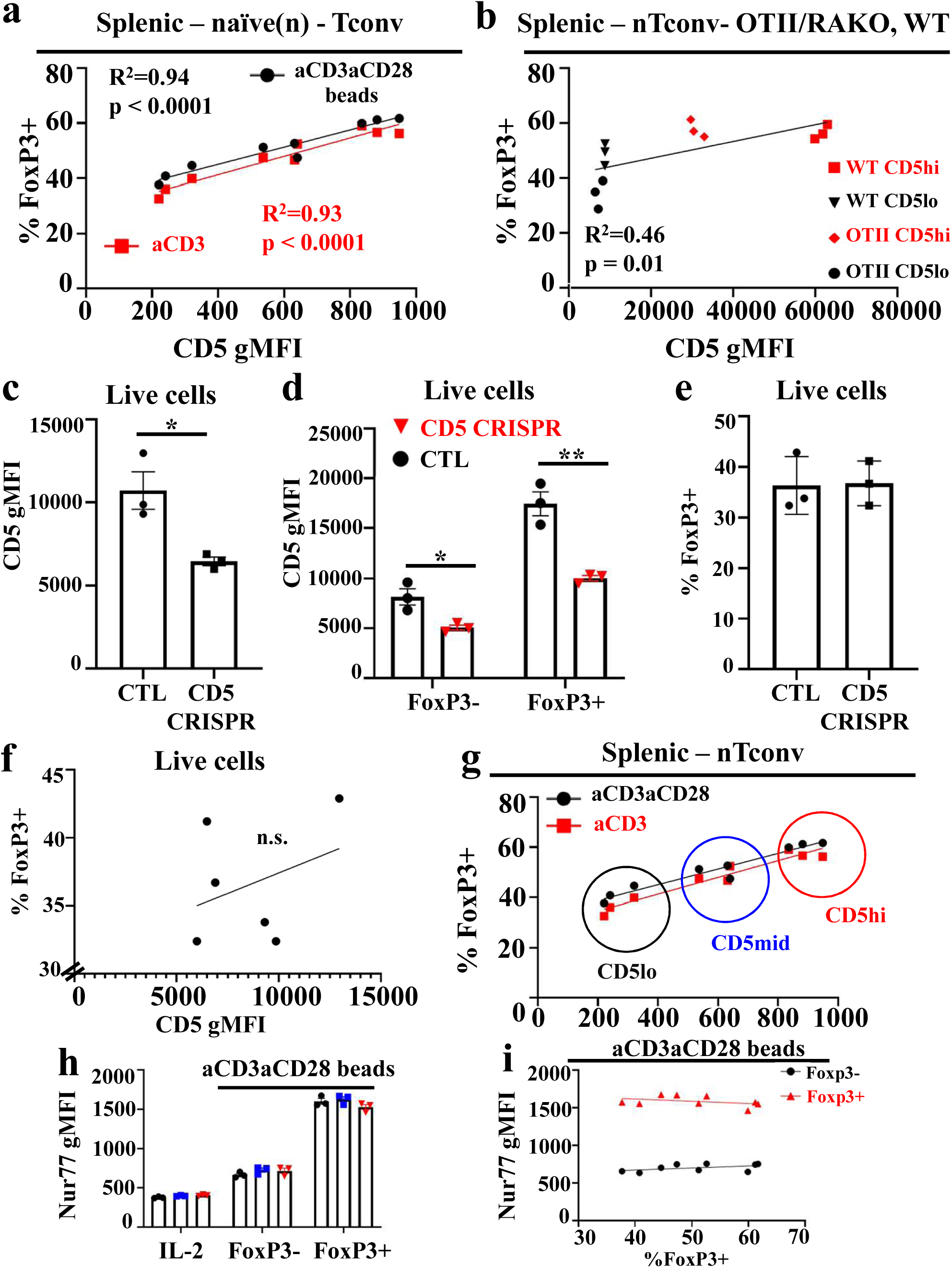
The CD5 molecule is not responsible for differences in iTreg induction. a. Correlation between initial sorted CD5 gMFI of naïve Tconv and their eventual %FoxP3+ after 3 days of in vitro culture: immobilized aCD3 (10ug/mL) or aCD3/CD28 beads (5ug/mL) with IL2 (50U/mL) and TGFb (2.5ng/mL). CD5lo and CD5hi cells were sorted from mice, in addition to an approximate ‘CD5mid’ population (n=3). Data representative of multiple experiments. b. Correlation between initial sorted CD5 gMFI of naïve CD5lo and CD5hi Tconv from WT and OTII/RAGKO cells and their eventual %Foxp3+ after 3 days of in vitro culture: soluble aCD3 (10ug/mL), IL2 (50U/mL), and TGFb (2.5ng/mL). Data representative of multiple experiments. c. CD5 gMFI of Tconv after 3 days of in vitro iTreg differentiation: aCD3/CD38 (2ug/mL), IL-2 (200U/mL), and TGFb (2ug/mL). Cultures either received guide RNA for CRISPR-mediated CD5 knockdown (CD5 CRISPR) or were transfection-only controls (CTL) (n=3). Data representative of multiple experiments. d. CD5 gMFI of Tconv on cultures from 3c, broken down into cells remaining Foxp3-or converting to Foxp3+. e. Percentage of cultures from 3c converting to FoxP3+. f. Correlation between final CD5 gMFI of cultures in 3c and %FoxP3+ conversion. g. Subjective grouping of cultures from 3a into CD5lo/mid/hi expressers for subsequent Nur77 comparisons. h. Comparison of day 3 Nur77 gMFI among cells from aCD3/CD28-stimulated cultures in 3g, including Foxp3-cells from cultures receiving IL-2 only (IL-2). i. Correlation of day 3 Nur77 gMFI in cells from aCD3/CD28-stimulated cultures in 3g with the percent conversion of cultures to Foxp3+.

These correlations were striking, however, interrogating CD5-related phenomena in T cells is not without its attendant pitfalls. Historically, studying the effects of CD5 in peripheral T cells has been complicated by the thymic effects of the molecule. Constitutive knockouts of CD5 impact T cell repertoire by modulating thresholds of negative selection (Tarakhovsky et al., 1995), resulting in concurrent changes in the repertoire and the proportion of thymic Tregs (Ordonez-Rueda et al., 2009). These studies established that the CD5 molecule is not necessary for Treg formation, at least in the thymus, but also that there would be potential artifacts in observed behavior from peripheral T cells developing in the absence of CD5; a hypothesis perhaps borne out from studies finding significant positive, neutral, and negative regulatory roles for the molecule, which in aggregate appear to be highly context-specific and limited in interpretation (Ceuppens and Baroja, 1986; McGuire et al., 2014; Perez-Villar et al., 1999).

Further complicating studies involving CD5 is that the ultimate function of the molecule remains a mystery, leaving open the possibility that CD5 itself is responsible for any observed differences in function. Therefore, given the extremely tight association between CD5 and iTreg induction, and previous work reporting direct CD5 functional influences on iTreg induction and T cell functionality, we felt it necessary to address to what extent the CD5 molecule itself may be functionally responsible for the heterogeneity in Foxp3+ differentiation among naïve peripheral T cells. To achieve this, we sought an experimental system whereby we could impair CD5 expression in naïve peripheral CD4 T cells only, without the thymic-selection-confounding complications associated with the use of T cells from constitutive CD5KO mice, or signaling inhibitors that likely have multiple cellular targets. To that end, we used an in vitro CRISPR-CAS9 system to impair expression of CD5 in naïve Tconv, which were subsequently incubated in our established iTreg conditions. The resulting cultures displayed a significant difference in surface CD5 levels by flow cytometry, establishing the efficacy of the CRISPR knockdown (Fig 3c). Further, the knockdown of CD5 was evident in both cells which had converted to Foxp3+ and those which had remained Foxp3-, suggesting there were no off-target effects of the RNA which might have caused cells to segregate into one phenotype or another (Fig 3d). Crucially, despite reduced CD5 expression, no alteration in the frequency of differentiated Foxp3+ T cells was observed between CTL and CD5- CRISPR Tcells (Fig 3e). To account for potential confounding variables in the delay of CD5 downregulation and signals necessary for Treg induction, we repeated the CRISPR experiments with a 24-hour rest of the cells between RNA nucleofection and the addition of iTreg induction reagents and found no difference in the results (data not shown). Finally, plotting the CD5 values by Foxp3+ induction percentage for all of the conditions and experimental replicates resulted in the disruption of the correlation between Foxp3 and CD5 expression (Fig 3f), establishing that while CD5 expression is coincident with Foxp3+ induction, CD5 functioning is not responsible for that correlation.

### CD5hi and CD5lo do not experience differential proximal TCR signaling

After ruling out differential functioning endowed by the CD5 molecule itself, we considered other factors that could influence acquisition of Foxp3 expression. In the thymus, it is known that levels of proximal TCR stimulation (along with co-stimulation) influence acquisition of the nTreg (Foxp3+) state (Jordan et al., 2001). Consistent with this, in our prior iTreg experiments (Fig 1), we observed that increasing concentrations of TCR agonist resulted in a concomitant increase in the frequency of Foxp3+ iTregs in both mature SP thymocytes and peripheral naïve CD4 splenocytes. Based upon these findings, we hypothesized that perhaps CD5hi peripheral nTconv cells are hardwired to perceive TCR stimulation more sensitively. To assess this possibility, we next measured Nur77 levels in our in vitro iTreg cultures.

It has been established that, like CD5, basal Nur77 levels are correlated with the strength of TCR signaling in thymic/self-antigen experience (Moran et al., 2011). Importantly, unlike CD5, Nur77 levels are both more transient and more rapidly upregulated in response to TCR stimulation, allowing us to observe potential differences in proximal TCR signaling ex vivo in our in vitro iTregs. In our cultures, we sorted Tconv across a range of CD5 values (Fig 3a), and then grouped the cultures into CD5 lo/mid/hi expressers and compared levels of Nur77 in response to stimulation (Fig 3g). In agreement with the established paradigm, upon TCR stimulation (either with [Fig 3h] or without CD28 [Supplemental Fig 2a]), we clearly observed increased Nur77 signals in all cells, with Foxp3+ cells expressing higher levels of Nur77 than Foxp3- cells. Surprisingly, however, there were no differences in Nur77 when comparing within either Foxp3+ or Foxp3 - cells between the CD5lo, CD5mid, or CD5hi cultures (Fig 3i, Supplemental Fig 2a). Consequently, we observed no linear correlation between Nur77 levels and %Foxp3 induction (Fig 3i, Supplemental Fig 2b). We therefore concluded that thymically imprinted predispositions coincident with naïve Tconv CD5 expression do not intrinsically alter subsequent perceptions of TCR stimuli.

### CD5hi iTregs more efficiently induce macrophage accumulation over PMNs

Before investigating further into intrinsic differences between CD5hi and CD5lo Tconv, we first wanted to determine if iTregs sourced from Tconv of varying self-affinities differed only in regard to their relative differentiation frequency, or if intrinsic differences in CD5 persisted beyond iTreg formation in a biologically relevant manner. Up to this point, we observed no difference in the outcome of our various in vitro experiments whether we utilized aCD3 stimulation or a combination of aCD3/aCD28 (Figs 1-3). Recent studies from our lab, however, determined that there is a distinct, qualitative difference between the types of iTregs created from these different culture systems, with aCD3 stimulation alone resulting in iTregs with the most stable expression of Foxp3 (Mikami et al., 2020). Based upon these findings, moving forward, unless otherwise noted, we elected to use only immobilized aCD3 stimulation for inducing iTregs in the remainder of the experiments.

In order test for functional differences between differentiated CD5hi and CD5lo Foxp3+ iTregs we employed a mouse model of airway allergy. In this model, mice are first sensitized to OVA through a series of injections of OVA with Aluminium Hydroxide (AlOH) (Ozkan et al., 2021). After sensitizing the mice, we adoptively transferred equal numbers of CD5lo or CD5hi iTregs into the mice, and then exposed the mice to aerosolized OVA for the next three days (Fig 4a). On the subsequent day, mice were euthanized and organs were collected for IHC and flow cytometric analysis. To evaluate antigen specific regulatory T cell activity, the transferred iTregs in these experiments were differentiated in vitro from CD5lo or CD5hi Tconv from OTII/RAGKO mice crossed with FoxP3-hCD2 reporter mice. This combination allowed us to compare, identify, and isolate ovalbumin specific, Foxp3+ iTregs by flow sorting for adoptive transfer.

**Figure 4.**
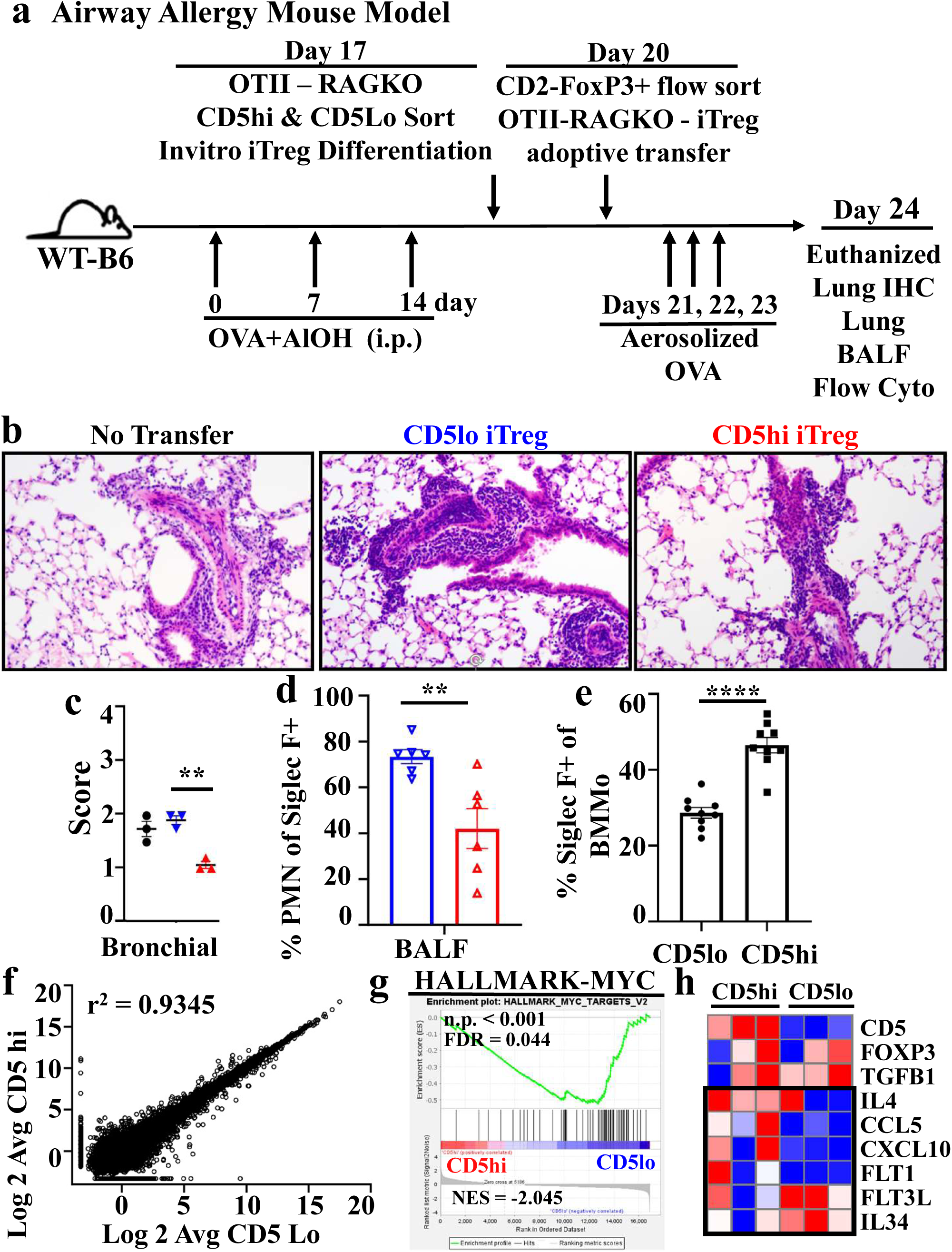
CD5hi iTregs more efficiently induce macrophage accumulation over PMNs. a. WT BL6 mice were sensitized to OVA with a series of i.p. injections over 2 weeks (OVA [400ug] and AlOH [1ug] in PBS, 200ul/mouse). iTregs were then induced from CD5- disparate Tconv of OTII/RAGKO/FoxP3-hCD2 mice and injected into OVA-sensitized mice, followed by 3 days of exposure to aerosolized OVA. Mice were sacrificed on day 24 and tissues were analyzed. b. Representative slides of lung HE staining, showing generally extensive pneumonitis among the groups. Color schema also used in figures 4c/d/e and Supplemental figure 2c (No transfer [black], CD5lo iTregs [blue], CD5hi iTregs [red]). c. Bronchial scoring of HE stained lungs (n=3). d. Fraction of Siglec F+ Bronchial Alveolar Lavage Fluid (BALF) cells that are PMN. Data combined from multiple experiments (n=6). e. Percentage of bone marrow monocyte (BMMo) cells Siglec F+ after 4 days coculture with CD5 disparate iTregs and soluble aCD3 (2ug/ml). Data combined from multiple experiments (n=9). f. Comparison of RNA expression profiles between iTregs from CD5lo/CD5hi Tconv, stimulated for 24hrs in vitro (2ug/mL soluble anti-CD3). Representative of multiple experiments. g. Gene set enrichment analysis (GSEA) on differentially expressed genes from sequence data in (g) and Myc-associated genes. h. Expression profiles of selected molecules on sequence data from (g).

Upon completion of the experiment, blinded clinical pathological scoring of H&E lung sections identified a significant depression in bronchial infiltration and inflammation in recipients of CD5hi iTregs (red triangles) compared to CD5lo iTreg recipients (blue triangles, Fig 4c). CD5lo iTreg recipients were indistinguishable from no transfer controls (Fig 4c. black circles). No differences between groups were observed in the alveolar or arteriole histological regions (Supplemental Fig 2c). Corroborating the histological bronchial observation, flow cytometric analysis of bronchial-alveolar lavage fluid clearly demonstrated morphologically distinct cell types collected from the airways of CD5hi iTreg vs CD5lo iTreg recipients (FSC/SSC, Supplemental Fig 2d). Flow cytometric analysis from BALF as well as single cell suspensions from dissociated lung tissues of iTreg recipients revealed both these populations to be CD45+ Siglec F+ positive populations, differing not only in their FSC/SSC properties but also F4/80+ expression (Supplemental Fig 2e). Based upon these characteristics, we ascribed these two populations to be PolyMorphoNuclear cells (PMN, Siglec F+ F4/80-, most likely Eosinophils) and macrophages (Macs, F4/80+ Siglec F+). In the BALF (Fig 4d), we observed a significantly higher ratio of PMN to macrophages from CD5lo iTreg recipients compared to CD5hi iTreg recipients. Collectively, these data support the hypothesis that even after differentiation into effectors, iTregs differentiated from CD5hi or CD5lo progenitors differ in their ability to influence the immune microenvironment in vivo.

Given the known interaction between Tregs and both monocyte/macrophages and PMN/granulocytes in the resolution of airway inflammation (Soroosh et al., 2013; Zhang et al., 2022), we hypothesized that factors secreted by the differentiated iTregs themselves might impact myeloid cell differentiation. To pursue this hypothesis, we performed in vitro bone marrow monocyte differentiation co-culture assays. In this assay, in vitro differentiated CD5hi or CD5lo iTregs were co-cultured with immature bone marrow monocytes from WT congenic mice under conditions of mild TCR stimulation. We observed that monocytes differentiated in the presence of CD5hi iTregs more frequently expressed Siglec F than those in the presence of CD5lo iTregs (Fig 4e). Interestingly, in vivo, Siglec F expression by macrophages and monocytes is associated with both migratory and alveolar macrophage phenotypes (Epelman et al., 2014; Scott et al., 2014). To our surprise, when we compared in detail the characteristics of Siglec F+ monocytes/macrophages derived from the CD5 disparate cocultures for markers of differentiation (Ly6G, CD64, F4/80) and activation (MHCII, CD80), we found no significant differences (Supplemental Fig 2f). This suggested to us that the changes induced in monocytes were less likely a factor of the CD5 disparate iTregs inducing distinct monocyte characteristics, and more likely reflective of a difference in relative abundance of similar molecules impacting the overall rate of monocyte differentiation.

To better determine potential differences between the effector functioning of these two groups of iTregs, we next performed RNASeq expression analysis on CD5hi- and CD5lo-derived iTregs after one day of TCR stimulation (as employed in our bone marrow differentiation assay). Upon plotting the average expression value for expressed genes, we observed a remarkable similarity between the populations by linear correlation (Fig 4f). We did find, however, that when we employed Gene Set Enrichment Analysis (GSEA) (Subramanian et al., 2005) to these data there were significant differences; notably in the gene sets ascribed to the transcription factor Myc (Fig 4g), IL-17 signaling, and eicosanoid metabolism (Supplemental Fig 3). While none of these pathways directly explained our in vitro or in vivo myeloid cell observations, they inspired us to examine RNA expression profiles specifically for cytokines and chemokines that might influence myeloid cell recruitment or differentiation (Fig 4h, black box). We observed that CD5hi derived iTregs had higher levels of IL-4 and CCL5, while CD5lo cells had higher levels of FTL3L and IL-34.

Mechanistically it is unclear how these differences in RNA expression would result in our in vitro and in vivo observations, but they do highlight that, while the CD5-disparate stimulated differentiated iTregs are quite similar, potentially important distinctions remain.

### CD5 coincident gene expression profiles start in the thymus and are retained through iTreg induction and activation

Thus far we have demonstrated that CD5-disparate naïve cells differentiate into Foxp3+ iTregs with different efficiencies in vitro (Fig 1) and in vivo (Fig 2). We also demonstrated that CD5-disparate Tconv which differentiate into iTregs in vitro have differential impacts on other immune cells, both in vitro and in vivo (Fig 4). We have eliminated both a direct role for CD5 and differential perception of TCR (as reported by Nur77 expression) (Fig 3) as accounting for these differences in iTreg differentiation. To further address potential differences in CD5hi and CD5lo naïve splenocytes which might account for iTreg induction (and to interrogate the possibility of an establishment or continuity of a CD5-linked T cell phenotype), we performed RNA expression profiling on CD5hi and CD5lo cells derived from naïve single-positive (CD4) thymocytes and naïve splenocytes, and also included iTregs derived from CD5hi and CD5lo naïve splenocytes. (Note that our previous RNA expression profiling (Fig 4) was performed on analogously differentiated iTregs, but only after re-stimulation with anti-CD3 [iTreg aCD3], as established for our co-culture differentiation system). Collectively, we have assembled a continuum of RNA expression profiles that track the difference between CD5hi and CD5lo cells from the mature SP thymocyte stage through iTreg differentiation and TCR re-stimulation.

Broad examination of our differential RNA expression results from CD5hi and CD5lo cells across the iTreg developmental spectrum (Fig 5a, Fig 4f) unsurprisingly revealed fewer distinct differences between the groups as cells progressed in their differentiation towards iTregs. To determine if the differences that existed between the CD5hi and CD5lo in each group persisted in the other developmental stages, we composed custom gene lists for each stage composed of genes (250) enriched in either CD5hi (Fig 5b) or CD5lo (Supplemental Fig3b), and then utilized these gene sets in the GSEA tool to perform pairwise comparisons of gene enrichment between all the developmental groups. Overall, we observed significant enrichment of the CD5hi genes derived from any one group in each of the other CD5hi groups. We observed a similar association within the CD5lo groups (Supplemental Fig 3b), with these enrichments being noticeably weakened only in the differentiated iTregs stimulated with anti-CD3 (far right column, bottom row). From these results, we inferred that a core group of genes may be co-regulated with CD5 in the thymus and persist from the thymic SP stage through iTreg differentiation. To determine if this was indeed the case, and to isolate which genes may be upregulated with CD5 in response to thymic self-antigen affinity, we composed an inferred regulatory network centered around CD5 by utilizing the 18 RNA Seq samples across the 3 different datasets (Thymus, Spleen, iTreg) and examining the Pearson linear correlation values between each expressed gene and CD5 (Fig 6a). After doing so, we were able to determine which genes positively or negatively varied in a tight relationship with CD5 (Fig 6b). Initial observations of the Top 25 genes positively associated with CD5 expression (Fig 6c) provided some validation of this approach, as genes previously established to be associated with differentiation or regulation of Treg identity or functionality (Izumo1r, ID2, TNFSF11) appeared toward the top--consistent with our observations that CD5hi cells harbor a predisposition for Treg differentiation (Francisconi et al., 2018; Miyazaki et al., 2014; Yamaguchi et al., 2007).

**Figure 5.**
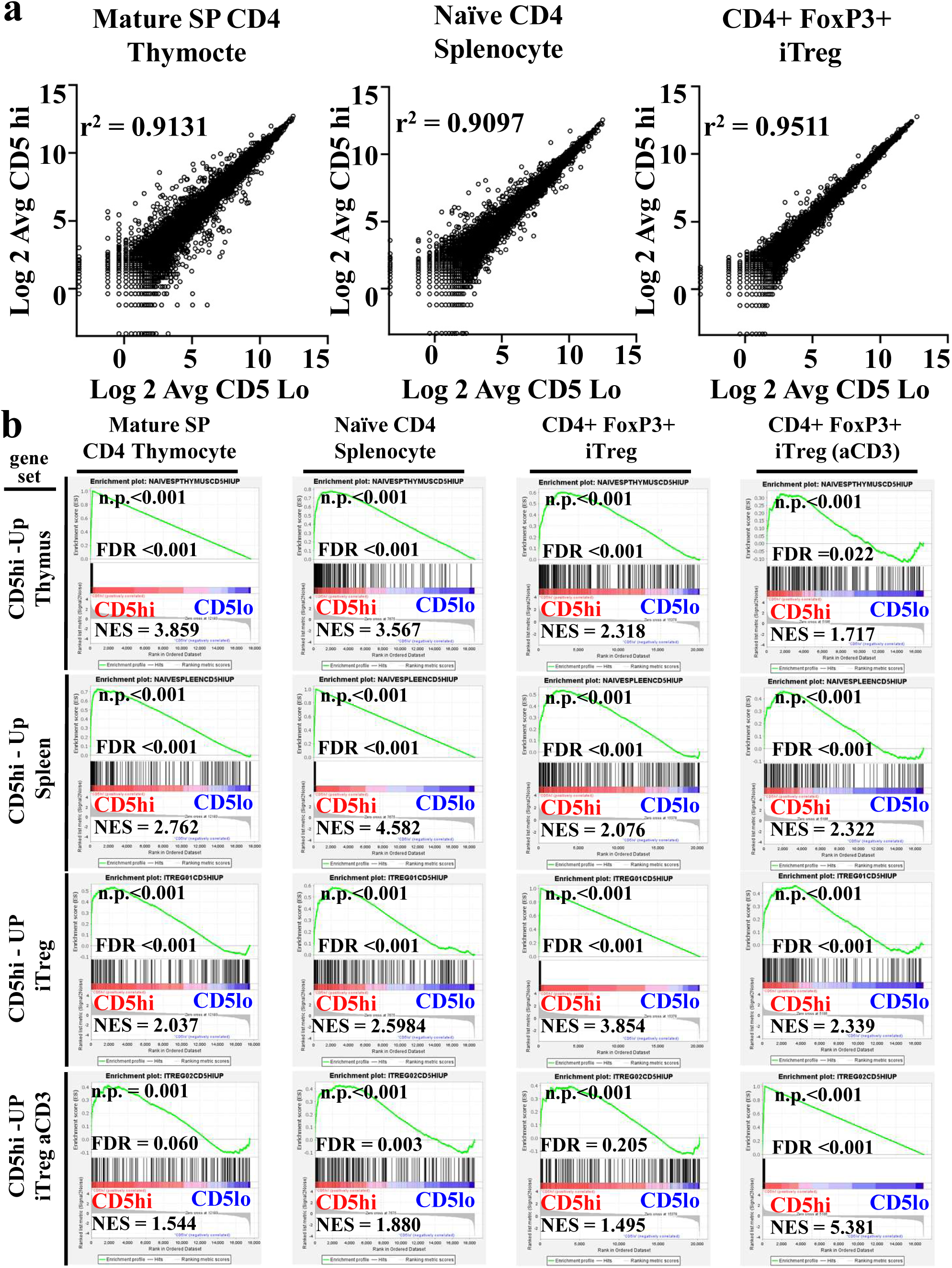
CD5 coincident gene expression profiles start in the thymus and are retained through iTreg induction and activation. a. Comparison of RNA expression profiles between CD5lo/CD5hi mSP thymocytes (left), Tconv (middle), and iTregs (right). Representative of multiple experiments. b. GSEA on top 250 differentially expressed genes from sequencing data in (a) and Figure 4g. Correlation of CD5hi gene signatures among groups.

**Figure 6.**
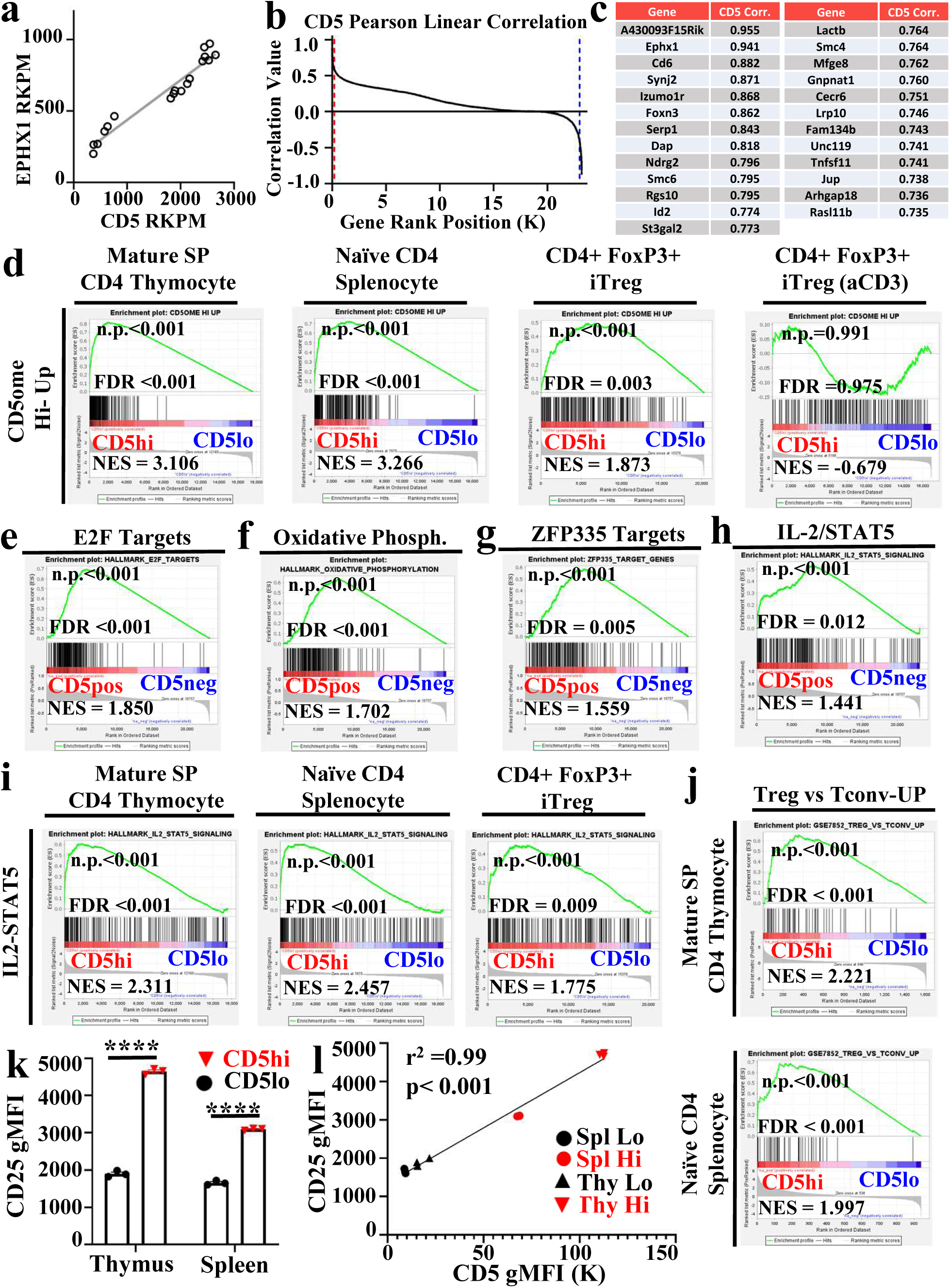
CD5ome associates with IL-2 signaling and predisposition for Treg identity. a. Correlation of CD5 and RNA expression with EPHX1 expression across spleen, thymic and iTreg populations b. Distribution of Pearson correlation values for all expressed genes with CD5. Red bar highlights 200 most positively correlated genes. Blue bar highlights 200 strongest negatively correlated genes. c. Top most CD5 positively correlated genes across RNA expression samples d. GSEA plot of top 200 positively correlated to CD5 genes in gene expression comparisons between CD5hi and CD5lo cells in CD4 single positive (SP) thymocytes, naïve splenocytes, iTregs and iTreg stimulated with aCD3 (left to right). Gene list preranked by gene pearson correlation value of gene expression to CD5 expression across all RNA samples subject to GSEA analysis using molecular signature database gene lists demonstrate CD5 expression is associated with; e. Cellular proliferation (E2F) f. Oxidative metabolism (Oxidative Phosphorylation) g. ZFP335 Targets h. IL-2/STAT5 signaling i. IL-2/STAT5 gene set enrichment applied to differential gene expression comparisons between CD5hi and CD5lo CD4 T cells as SP thymocytes, splenocytes and iTregs j. GSEA enrichment plot of Treg vs Tconv upregulated gene set in CD4 thymocytes (top) and splenocytes (bottom) k. CD25RNA expression values in CD5hi (red) and CD5lo (black) thymus and splenocyte derived CD4 T cells. (n=3). Data representative of multiple experiments. l. Correlation between CD5 and CD25 expression in groups from (k).

We next composed gene sets from this inferred CD5 transcriptional expression network (CD5ome) from the top 200 (Fig 6b, red line) and bottom 200 (Fig 6b, blue line) most correlated genes, and repeated the GSEA analysis to determine the presence and persistence of these genes in comparisons of CD5hi vs CD5lo across the developmental spectrum (Fig 6d). We again saw a strong correlation of these de novo CD5 associated genes with CD5hi cells, from mature thymocytes through iTreg differentiation (though the relationship weakens under aCD3 stimulation of iTregs). Overall, the RNA expression is consistent with the hypothesis of a preset thymic program correlating with CD5 expression that persists through iTreg differentiation.

### CD5hi Tconv have an increased sensitivity to IL-2

To determine which genes or pathways might be incorporated in this CD5 associated transcriptome, we performed GSEA analysis on this CD5 correlation ranked list utilizing canonical gene sets from the molecular signature database. We observed significant enrichments in multiple gene sets associated with cell cycle and proliferation (e.g. E2F, Fig 6e) and metabolism (e.g. oxidative phosphorylation [Fig 6f]). We also observed enrichment in genes associated with the transcription factor zinc finger protein 335 (ZFP355, Fig 6g) recently ascribed to have a role in thymic development and regulation of the genes BCL6 and RORC (Wang et al., 2022). Finally, we observed enrichment of genes associated with IL-2 and STAT5 signaling (Fig 6h). This IL-2/STAT5 enrichment was notable, as it appeared in our CD5hi vs CD5lo GSEA analyses of each of the independent cell groups as well (thymocyte, splenocyte, iTreg, Fig 6i). Further corroborating these associations between IL-2, CD5, and thymic imprinting toward iTreg identity, our GSEA analyses also revealed a significant Treg signature when comparing CD5hi to CD5lo cells from the mature SP thymocyte and naïve CD5 splenocyte populations (Fig 6j). Given the reported centrality of STAT5 signaling for anchoring the stability and functionality of the Foxp3 transcription factor in regulatory T cells (Burchill et al., 2007; Chen et al., 2011), we chose to examine the association of IL-2 signaling with CD5 levels as an explanation for the elevated proclivity of CD5hi to become iTreg.

The most logical explanation for increased IL-2 signaling between CD5hi and CD5lo cells in the same environment would be the presence of increased IL-2 receptor (IL-2R) components on CD5hi cells. Utilizing flow cytometry, we determined the expression levels of the IL-2Ra (CD25) and IL-2Rb (CD122) components. Consistent with our RNA expression and GSEA results, our flow cytometric analyses found a significant increase in the expression of CD25 on CD5hi compared to CD5lo in T conv from both the thymus and spleen (Fig 6k), and revealed a tight association between CD25 and induced Treg frequency (Fig 6l). No such patterns were found for IL-2Rb (Supplemental Fig 4c).

Next, we checked to see if these observed differences in receptor expression translated into differential responsivity to IL-2. To accomplish this, we treated CD5hi and CD5lo nTconv splenocytes with IL-2 and used intracellular phosphoflow cytometry to observe the phosphorylation state of STAT5, the known downstream target of IL-2R activity. CD5hi nTconv displayed a clear response to IL-2, whereas the CD5lo cells showed none (Fig 7a). It was noted that, while significant, these responses were muted compared to those observed when nTregs were treated with IL-2 (supplemental Fig 4d). Previously, Surh and Sprent reported that CD5hi expression was associated broadly with an increased sensitivity to cytokines, but were interested primarily in naïve CD8 T cells (Cho et al., 2010). Thus, we repeated these studies with IL-4, IL-6, and IL-7 to determine the specificity of our observations for IL-2 and STAT5 signaling. Surprisingly, yet consistent with the observations of Surh and Sprent, we noted increased pSTAT6/3/5 signaling, respectively, in response to stimulation with these cytokines (Fig. 7b). This was surprising simply because pathways associated with these cytokines did not appear among those upregulated between CD5hi and CD5lo in our RNA expression analysis. Surh and Sprent attributed their observed differential cytokine signaling among naïve CD8 T cells to an increase in the lipid raft composition of CD5hi cells, which thereby conferred more efficient cytokine signaling. Correspondingly, we stained our CD5lo and CD5hi cells with fluorescently labeled cholera toxin for the quantification of lipid rafts and, in agreement with the observations of Surh and Sprent, determined that CD5hi cells had higher amounts of lipid rafts (Fig 7c). Collectively, these data suggest that CD5hi cells have dual advantages over CD5lo cells in responding to IL-2: first, they express higher amounts of the high affinity IL-2 receptor, and second, through lipid raft aggregation, they are likely able to make greater use of limited IL-2 signaling, setting the stage for increased iTreg differentiation.

**Figure 7.**
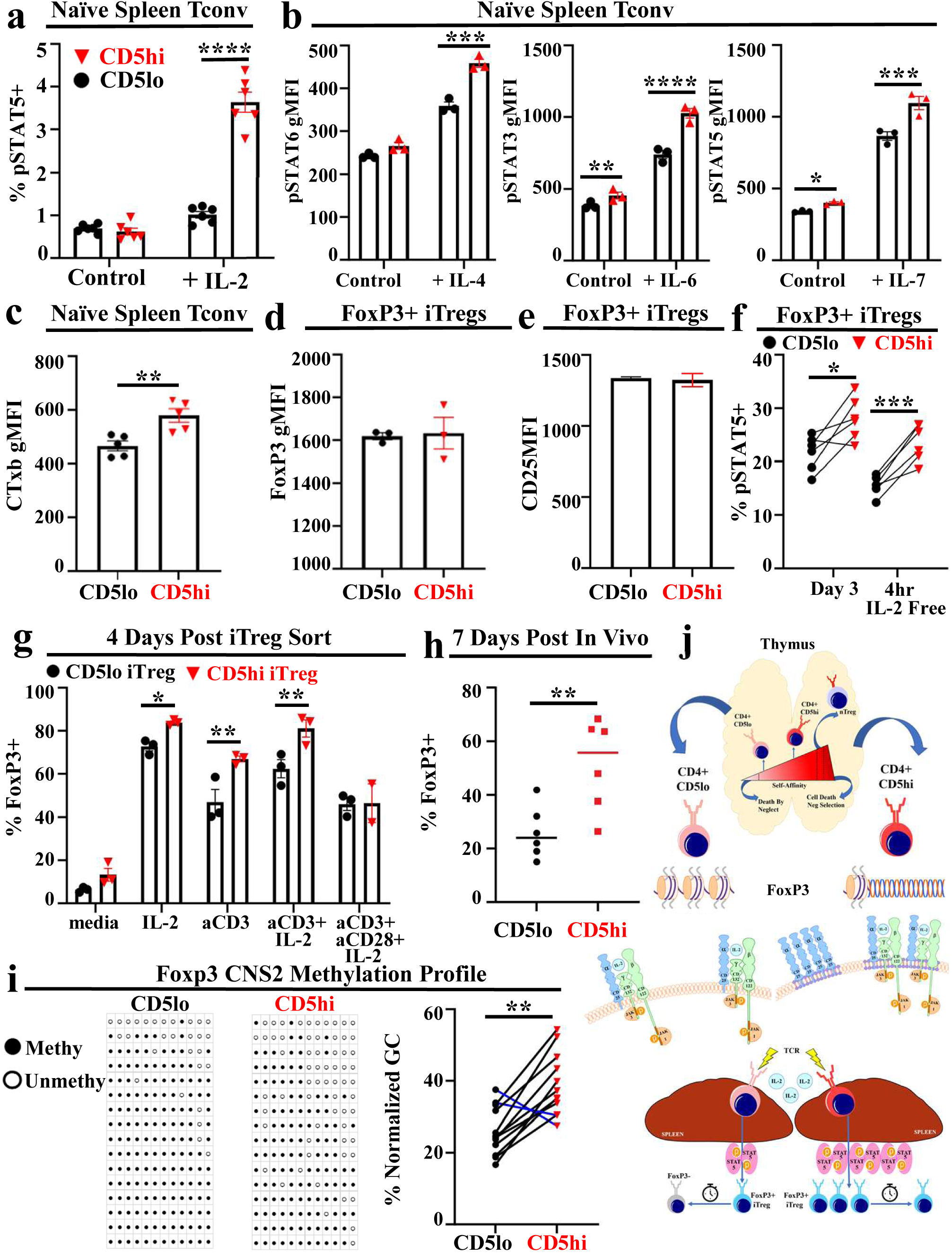
CD5 differences persist in the effector stages of iTregs and impact stability of Foxp3 expression. a. Percentage of sorted Tconv cells staining positive for phospho-STAT5 with or without 30 minutes of IL-2 stimulation (n=6). Data combined from two experiments. b. Percentage of sorted Tconv cells staining positive for phospho-STAT proteins after cytokine stimulation: IL4 (left); IL6 (middle); IL7 (right) (n=3). Data representative of multiple experiments. c. Comparison of lipid raft levels in naïve Tconv, as evidenced by cholera toxin b (CTxb) (n=5). Data combined from two experiments. d/e. Comparison of Foxp3 (d) and CD25 (e) protein levels in iTregs from differently sourced Tconv (n=3). Data representative of multiple experiments. f. Percentage of iTregs staining positive for phospho-STAT5, either at the end of a 3 day in vitro differentiation assay, or an additional 4 hours in the absence of IL2 (n=6). Data are combined from two experiments. g. Percentage of sorted iTregs staining positive for FoxP3 expression 4 days after differentiation/re-plating: IL2 (50U/mL), soluble aCD3 (0.5ug/mL), aCD3/CD28 beads (1.0ug/mL) (n=3). Data representative of multiple experiments. h. Percentage of in vitro induced iTregs staining positive for Foxp3 7 days after adoptive transfer into normal, lymophoreplete mice (n=6). Data combined from two experiments. i. Representative methylation pattern of Foxp3 CNS2 region from CD5lo or CD5hi iTregs (left), paired comparison of the different CNS methylation sites between the two groups of iTregs (right) (n=6). Data combined from two experiments. j. Model slide of experimental results.

### CD5 differences persist in the effector stages of iTregs and impact stability of Foxp3 expression

The question still remains how these IL-2 signaling differences translate into the effector, differentiated iTreg stage, and whether these differences impact iTreg functionality. Surprisingly, when run in a standard in vitro suppression assay, CD5lo iTregs showed no deficit in inhibiting the proliferation of Tconv cells compared to CD5hi Tregs (data not shown). Crucially, because these cells were isolated based upon their CD2 reported Foxp3 expression (Supplemental Fig 4e), at the initiation of the assay, iTregs from both CD5hi and CD5lo cells were known to express robust and equivalent levels of Foxp3 (Fig 7d); an important indicator of Treg functionality. It was previously demonstrated that IL-2/STAT5 signals are neither necessary nor sufficient for acquisition of Foxp3 expression, however, over-expression of STAT5 during differentiation does result in a greater yield of Foxp3+ cells at the end of culture (Burchill et al., 2007). It has subsequently been repeatedly demonstrated that IL-2 signaling influences chromatin remodeling of the Foxp3+ locus, enhancing Treg identity and stability (Ohkura and Sakaguchi, 2020). To determine if Foxp3 instability and chromatin remodeling are behind the differences observed between CD5hi and CD5lo cells, we again differentiated iTregs from CD5hi or CD5lo cells from Foxp3-hCD2 mice (Supplemental Fig 4e) and monitored the cells for stability in Foxp3 expression of differentiated and sorted iTregs. We also assessed which of our previous observations of differences between CD5hi and CD5lo splenic and thymic Tconv cells carried over into the effector stage of iTreg by examining some of these specific characteristics in fully differentiated iTregs. First, we assessed CD25 expression in differentiated iTregs on day of isolation by flow cytometry and, unlike the case for splenocytes and thymic naïve cells, we found no difference between iTregs derived from CD5lo and CD5hi progenitors (Fig 7e). We did, however, observe that CD5hi-derived iTregs demonstrated higher levels of pSTAT5 signaling than CD5lo cells, both at the conclusion of 3 days of differentiation, and when re-cultured in media lacking IL-2 (Fig 7f), reminiscent of the suppressed pSTAT5 observed in splenic and thymic CD5lo T conv.

While interrogating differences between CD5hi- and CD5lo-derived iTreg cells for Foxp3 stability, we observed that as soon as 4 days after sorting the differentiated iTregs there were statistically significant drops in Foxp3 expression within the CD5lo derived iTregs compared to CD5hi derived iTregs across a variety of in vitro culture conditions, including conditions supplemented with 50U/mL of IL-2 (Fig 7g). Curiously, even increasing the IL2 concentration of the culture to 8x the normal levels had no effect on rescuing/maintaining the Foxp3+ phenotype (Supplemental Fig 4f). Throughout these assays we observed no significant levels of cell death or disparate levels of cell death that could account for this drop in Foxp3 expression. Mirroring this response, when congenically disparate CD5hi- or CD5lo-derived Foxp3-hCD2 iTregs were re-isolated 7 days after being injected in WT hosts, the frequency of CD5lo cells expressing Foxp3 dropped significantly compared to Tregs derived from CD5hi cells (Fig 7h). As a final survey into the stability of Foxp3 in the two different groups, we compared the methylation of the Foxp3 CNS2 region in iTregs directly after their induction in vitro (Ohkura and Sakaguchi, 2020). We observed that, even at this time point (where Foxp3 expression is equal and IL-2 is provided), CD5hi cells demonstrate a greater level of demethylation at 10 of the 12 CNS2 methylation sites compared to CD5lo cells; indicative of greater gene accessibility and stability (Fig 7i), and predictive as well as consistent with the fate we observed of Foxp3 in analogous cells in vitro and in vivo.

## Discussion

A high affinity for self has long been associated with Tregs, especially (and most explicitly) in the relationship between highly self-reactive CD4+ thymocytes and their proclivity for developing into thymic Tregs (nTregs). This clear association becomes weaker in the periphery, with recent studies examining the relationship between self-affinity (however it is defined) and iTreg/pTreg formation pointing towards a likely correlation, but differing on some of the finer points. In an attempt to unify the field on this point, we undertook the above experiments, and found a robust correlation between self-affinity (as defined by CD5) and proclivity for iTreg induction in vitro, across different stimulatory conditions and T cell repertoires. This finding also extended to in vivo pTreg formation, even when conducted in the non-lymphopenic environment of physiologically normal mice. These data were further supported by the RNAseq/GSEA experiments, which showed a strong significance to the correlation between the gene signature of CD5hi cells, at both the mature-single positive and peripheral naïve stages, and that of Tregs, suggesting that these cells are also genetically poised to become iTreg/pTreg.

An outstanding question in any study of T cell self-affinity is the effect of the CD5 molecule itself on the phenomena under investigation. With the important caveat of a very intriguing recent study into the importance of CD5 in T cell-DC interactions (He et al., 2023), the molecule proves surprisingly resilient to close inspection--likely due to the difficulty in teasing apart its effects in the thymus during development and in peripheral functioning. Thus, when stratifying T cells along CD5 and observing subsequent functional differences between the cells, it is tempting to ascribe those differences to the as-yet full undefined functioning of the molecule itself. Controlling for this potential pitfall is complicated, as complete knockouts of CD5 result in an altered T cell development (as evident in the altered T cell repertoire), which raises the possibility that developing in the absence of CD5 will result in artifactual changes to normal T cell functioning, thereby interfering with subsequent observations. Of course, the creation by the Klechevsky group of a CD5 conditional knockout mouse in the study mentioned above (and similar efforts underway in this lab) have largely solved this problem going forward, but, for our current studies, we controlled for the potential involvement of CD5 with a CRISPR-Cas in vitro knockdown system, and were able to determine that downregulation of expression of the CD5 molecule did not influence rate of iTreg differentiation post the single positive thymic maturation stage.

A further unstudied aspect of T cell self-affinity and its effect on cell fate and functioning is the degree to which this phenomenon extends into the secondary/effector phase of the cell. Previous studies into this particular question do exist, but have largely considered the effector cells as a unified whole—perhaps most elegantly by the Allen group in their study determining which of two monoclonal TCR T cells, of identical affinity for cognate bacterial antigen but differing CD5 values, displayed better effector functioning (Milam and Allen, 2015; Persaud et al., 2014). To approach this concept from a different angle, we wanted to determine, on a per-cell basis, if T cells maintained some aspect of the heterogeneity displayed during the naïve stage into the effector phase. Using our Foxp3-reporter mice, we observed that there are, indeed, a number of intriguing differences between the iTregs sourced from Tconv of varying self-affinities, with the most striking difference being the significant difference in responsivity to IL-2 and the relative stability of Foxp3 expression between the two groups. Indeed, some of the other observed differences between the two iTregs, such as the difference in induced differentiation of the BMMo into Siglec F+ cells, could potentially be explained by this tendency of the CD5lo iTregs to quickly lose their ‘Treg-ness’. There remain some intriguing differences in the RNAseq data between the two groups of iTregs, however, which suggest there may be further differences in effector function that can be further teased out through additional, thorough studies. Ultimately this study further supports a model whereby thymic self-antigen exposure predetermines cellular attributes that carry over into the naïve repertoire, impacting the functional effector status of subsequently differentiated cells. This has obvious implications for both adoptive and chimeric antigen receptor, cytotoxic, and regulatory T cell therapy. Identifying cells derived from strong thymic self-peptide-MHC interactions, by CD5 or some other means, could lead to greater stability of cellular products and therefore greater clinical efficacy. Our report joins a growing number of manuscripts which implicate a role for original thymic antigen experience (original thymic antigenic sin) as a significant contributor to shaping subsequent immune responses.

## EXPERIMENTAL MODEL AND SUBJECT DETAILS

### Mice

All animals used in this study were maintained in specific pathogen-free conditions. All experiments using animals were performed following the guidelines on animal welfare of Osaka University.

C57BL/6 mice were purchased from CLEA Japan, inc. C57BL/6 (CD45.1) congenic and OTII/RAGKO mice were bred in our animal facility. Foxp3-IRES-DTR-GFP mice (FDG) and Foxp3-IRES-hCD2-hCD52 mice (Foxp3-hCD2) were previously described (Kim et al., 2007; Komatsu et al., 2009). Experiments were performed using male or female mice aged 7 to 12 weeks old.

## METHOD DETAILS

### Antibodies and reagents

Antibodies, reagents and critical commercial assays used in this study are listed in the Key resource table.

### Flow cytometry analysis and cell sorting

For cell surface staining, prepared cell suspension was incubated with fluorescence-conjugated antibody cocktail in flow cytometry (FACS) buffer (2% FCS and 1mM EDTA in PBS or RPMI1640) for 20-40 min at 4C. Live/dead cell staining (ThermoFisher) was added if necessary. For intracellular staining, we used the Foxp3 staining buffer kit (eBioScience) according to manufacturer’s instruction. For lipid raft staining, we used the lipid raft staining kit (Molecular Probes) according to manufacturer’s instruction.

Flow cytometry analysis was performed using an LSRFortessa (BD). Cell sorting was performed using FACSAriaIII or FACSAriaFusion (BD). For cell sorting of splenocytes and lymphocytes, pre-enrichment of CD4+ cells was performed using a CD4+ isolation kit (Miltenyi) according to manufacturer’s instruction.

Gating strategies for the cell fractions used in this study:

thymic mature single-positive cells: CD4+, CD8-, B220-, Foxp3-eGFP-OR Foxp3-hCD2-, CD44lo, CD25-, MHCI+, CD5lo (15%) or CD5hi (15%)
naïve splenocytes: CD4+, CD8-, B220-, Foxp3-eGFP-OR Foxp3-hCD2-, CD44lo, CD25-, CD5lo (15%) or CD5hi (15%)

### Phospho-staining

T cells of interest were incubated for 30 minutes with cytokines at the concentration noted in figure legends, after which they were washed once with PBS. Cells were then fixed for 10 minutes at 37C using pre-warmed Lyse/Fix buffer (BD), after which they were washed once with chilled FACS buffer. Prechilled Perm buffer III (BD) was then added to cells, which were incubated on ice for 30 minutes, and subsequently washed twice with prechilled FACS buffer. Cells were then resuspended in chilled FACS buffer containing the phospho-antibody of interest, and incubated overnight at 4C, followed by flow cytometry analysis.

### Cell culture, Treg cell induction, and in vitro iTreg cell stability assay

Purified T cells (purity > 95%) were cultured in vitro in a 37C, 5% CO2 incubator using RPMI1640 culture medium supplemented with 10% FCS (v/v), 60 mg/ml penicillin G, 100 mg/ml streptomycin, and 0.1 mM 2-mercaptoethanol. For iTreg cell induction from naïve thymocytes and splenocytes, the standard procedure was 1×10^5^ sorted T cells stimulated in a flat-bottom 96-well plate using plate-coated anti-CD3 antibody (10ug/mL) in the presence of 50 U/ml IL-2 and 2.5 ng/ml TGF-b. Other experiments using varying amounts of reagents, and also CD3/CD28 stimulatory beads (Gibco), list the final concentrations in figure legends. For analysis of FoxP3+ stability, iTregs (hCD2+/dead cell-) were sorted from in vitro cultures at differentiation day 3, and 2.0×10^4 - 5.0×10^4 cells were plated in flat-bottom plates in culture conditions noted in figure legends.

### Bone marrow monocyte differentiation

Bone marrow cells were prepared from femur and tibia of CD45.1+ wild-type C57BL/6 mice using 23G needle and syringe. Red-cell lysis was performed (Sigma) followed by monocyte isolation by kit (Miltenyi) according to manufacturer’s instruction. 5.0 x 10^4 of the resulting monocytes were subsequently plated in flat-bottom 96-well plates, co-cultured with equal numbers of iTregs sourced from CD5lo or CD5hi Tconv with 2ug/mL soluble anti-CD3. At day 4, cells were rinsed with cold PBS, adherent cells were detached with a scraper, and analysis was performed.

### In vitro CRISPR/Cas9

Peripheral naïve Tconv were isolated from the splenocytes of unmanipulated mice (2.0 x 10^6 cells/condition). RNA oligo (CD5 gRNA + tracrRNA) and RNP complex (gRNA/tracrRNA + Cas9) formation were completed according to manufacturer’s instructions (IDT). Subsequent nucleofection was done according to manufacturer’s instruction (Lonza), and was accomplished using the 4-D Nucleofection, X-Unit (Lonza, program: unstimulated mouse T cells). Cells were washed immediately after nucleofection to remove nucleofection reagents and improve survival.

### RNA sequencing

RNA was isolated using an RNeasy kit (Qiagen) according to manufacturer’s instructions. RNA libraries for sequencing were either prepared from samples utilizing Ion Express Plus fragment library kit (Thermo Fisher Scientific) and sequenced on an Ion S5 sequencer; or they were prepared using KAPA Hyper plus library kit (Roche) and sequenced on DNBSEQ (MGI) and NovaSeq (Illumina) sequencers, using 100bp paired end and 50bp single end reads respectively. All Sequences were preprocessed with Trimmomatic (version 0.39) and redundantly mapped against the mouse reference genome (GRCm38) using hisat2 (version 2.1.0) and bowties2 (version 2.3.5). Mapped reads were converted to BAM files by samtools (version 1.9) and PCR duplicate reads removed by picard MarkDuplicates (version 2.20.5-SNAPSHOT). Normalized counts were calculated by htseq (0.11.2), FPKM counts by cuffnorm (version 2.2.1) and tsv output files generated using ikra v1.2.3(https://github.com/yyoshiaki/ikra).

### Gene Set Enrichment Analysis (GSEA) and inferred CD5 regulon

For GSEA analysis, FPKM normalized gene counts for 3 samples from CD5hi and CD5lo cells, from either T conventional splenocytes, T conventional thymocytes, iTreg or iTregs post stimulation were inputed into the GSEA tool (GSEA 4.3.2 to identify enriched gene sets from the Molecular Signature Database (MSigDB v 2023.1, Hallmark, M2-Canonical Pathways & GTRD). GSEA tool was then used to rank differentially expressed genes and generate custom genes sets for each CD5hi and CD5lo group which were subsequently compared across groups. Custom gene lists for each stage were composed of genes (250) enriched in either CD5hi (Fig 5b) or CD5lo groups. The inferred CD5 regulatory network was composed by using expression values from 18 RNA Seq samples across the 3 different datasets (Thymus, Spleen, iTreg) and examining the Pearson linear correlation values between each expressed gene value and CD5. We next composed gene sets from the top 200 most positively correlated and bottom 200 (most negatively correlated) genes, and repeated the GSEA analysis to determine the presence and persistence of these genes in comparisons of CD5hi vs CD5lo across all CD5hi vs CD5lo datasets.

### Airway allergy induction

Wild-type C57/BL/6 mice were injected i.p. with a solution of 400ug OVA (Serva) and 1ug AlOH (Sigma) in 200ul PBS at experiment days 0, 7, and 14. On day 17, splenic Tconv from OTII/RAGKO mice were sorted into CD5lo and CD5hi populations, and cultured in in vitro Treg differentiation conditions. At day 20, iTregs were sorted from the in vitro cultures (huCD2+/dead cell-), and adoptively transferred into OVA-sensitized WT mice (1.0 x 10^5 iTreg/mouse). On days 21, 22, and 23, OVA-sensitized mice were consolidated into an air-flow restricted cage, and a 400ug/mL solution of OVA (Serva) in PBS was nebulized (Omron) and pumped into the cage for an hour. At day 24, mice were euthanized, lungs were perfused with PBS, BALF was collected, and the lungs were split in two, with half being used for flow analysis, and the other half placed in 10% formaldehyde for tissue histology.

### Tissue histology analysis

Freshly-isolated tissues were immediately fixed by 10% formaldehyde. H&E staining and microscopy slide preparation was performed by the Lab Solution Co. of Osaka. For scoring of lung inflammation, the alveolar space, bronchioles, and arterioles were scored separately on a scale of 0: no inflammation, 1: mild inflammation, 2: mild inflammation with notable clustering, 3: moderate inflammation with large clustering and without significant tissue destruction, 4: large diffuse or large clustering of inflammation with destruction of tissue.

### DNA methylation analysis

Bisulfite sequencing analysis of the Foxp3 locus was performed according to a previously established and reported protocol (Arai et al., 2022).

---

**Supplemental Figure 1.**
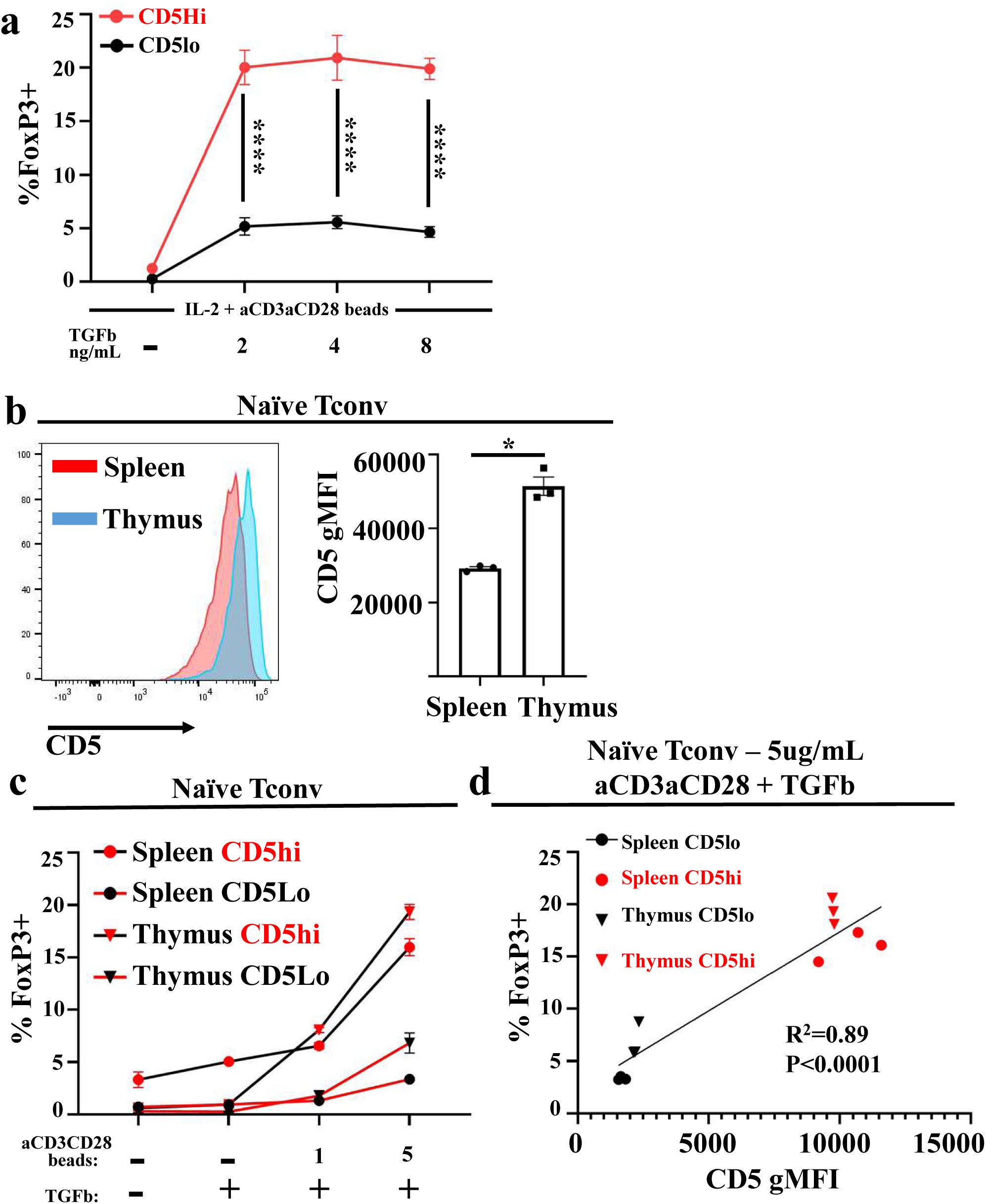
a. Percentage of iTreg obtained from in vitro induction experiments on CD5-disparate naïve Tconv splenocytes: aCD3/CD28 beads (5ug/mL) with IL2 (50U/mL) and varying amounts of TGFb for 3 days (n=3). Data representative of multiple experiments. b. Representative CD5 expression on CD4+ naïve splenocytes and mature single-positive CD4+ thymocytes (left), comparison of CD5 gMFI between the two populations (right, n=3). Data representative of multiple experiments. c. Percentage of iTreg obtained from in vitro induction experiments on CD5-disparate naïve Tconv splenocytes and mature single-positive thymocytes: varying amounts of aCD3/CD28 beads (ug/mL) with IL-2 (50U/mL) and TGFb (2.5ng/mL) for 3 days (n=3). Data from c (aCD3/CD28 5ug/mL); %FoxP3+ plotted against original sorted CD5 gMFI of various cultures.

**Supplemental Figure 2.**
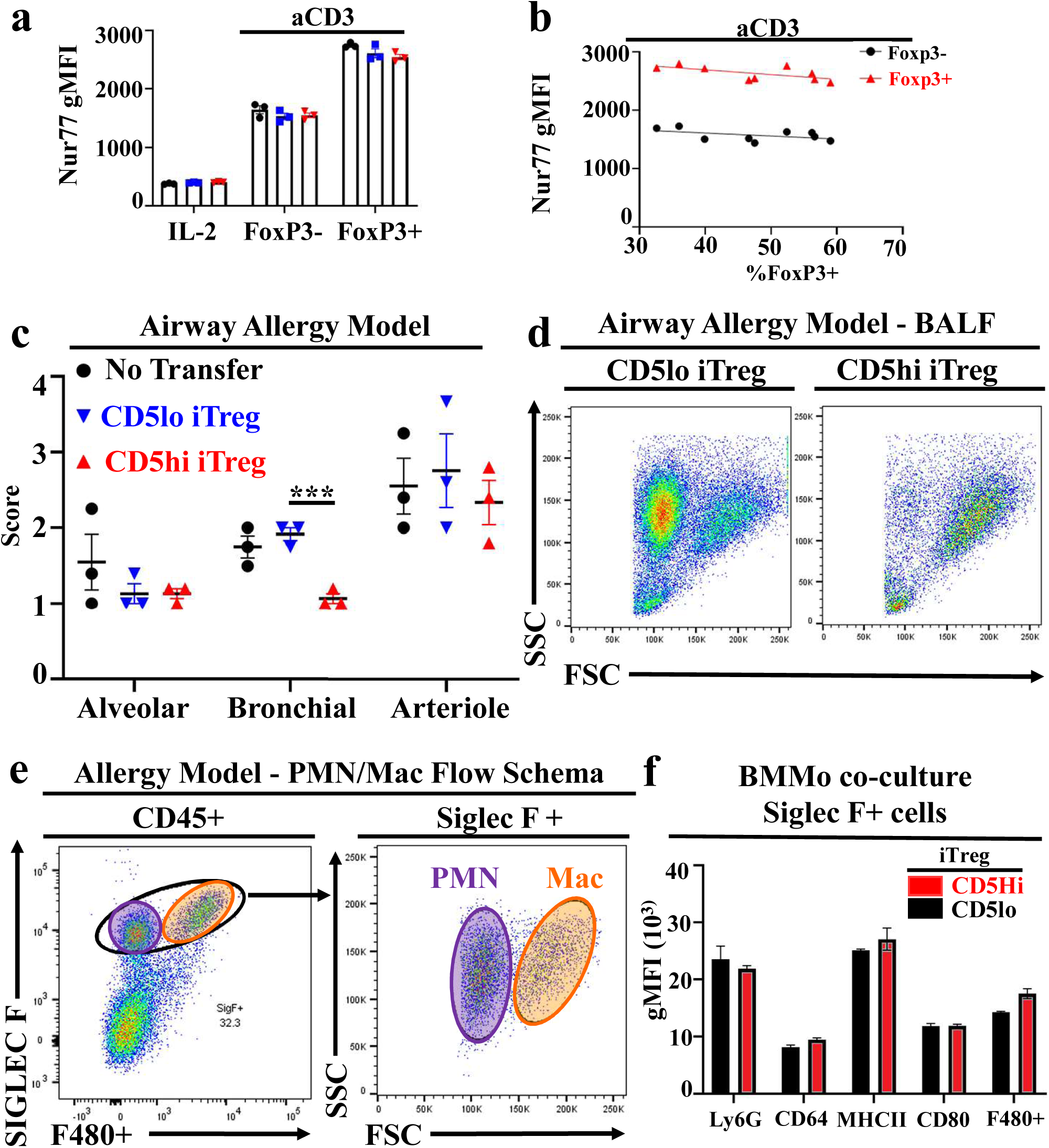
a. Comparison of day 3 Nur77 gMFI among cells from immobilized aCD3-stimulated cultures in 3g, including FoxP3- cells from cultures receiving IL-2 only (IL-2). b. Correlation of day 3 Nur77 gMFI in cells from immobilized aCD3-stimulated cultures in 3g with the percent conversion of cultures to Foxp3+. c. Full pneumonitis scores from figure 4c. d. Representative forward/side scatterplots of BALF fluid from experimental airway allergy mice receiving adoptive transfer of iTregs. e. Flow schema for differentiating between PMN and macrophages in experimental samples. Comparison of gMFI levels of various surface markers in SiglecF+ cells from BMMo differentiation cultures day 4 (n=3). Data representative of multiple experiments.

**Suppplemental Figure 3.**
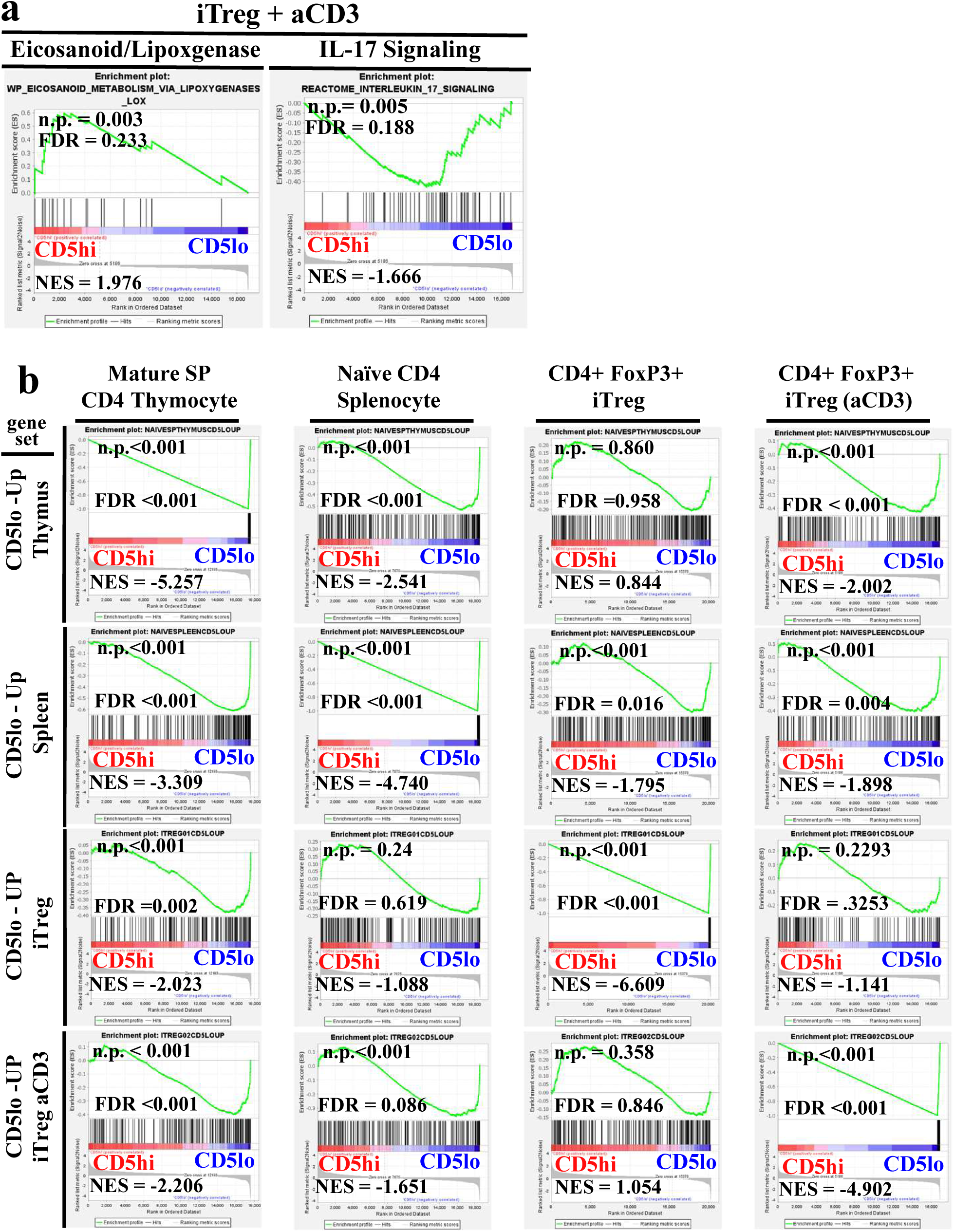
a. GSEA on differentially expressed genes from sequence data in Figure 4g. b. GSEA on top 250 differentially expressed genes from sequencing data in (a) and Figure 4f. Correlation of CD5lo gene signatures among groups.

**Supplemental Figure 4.**
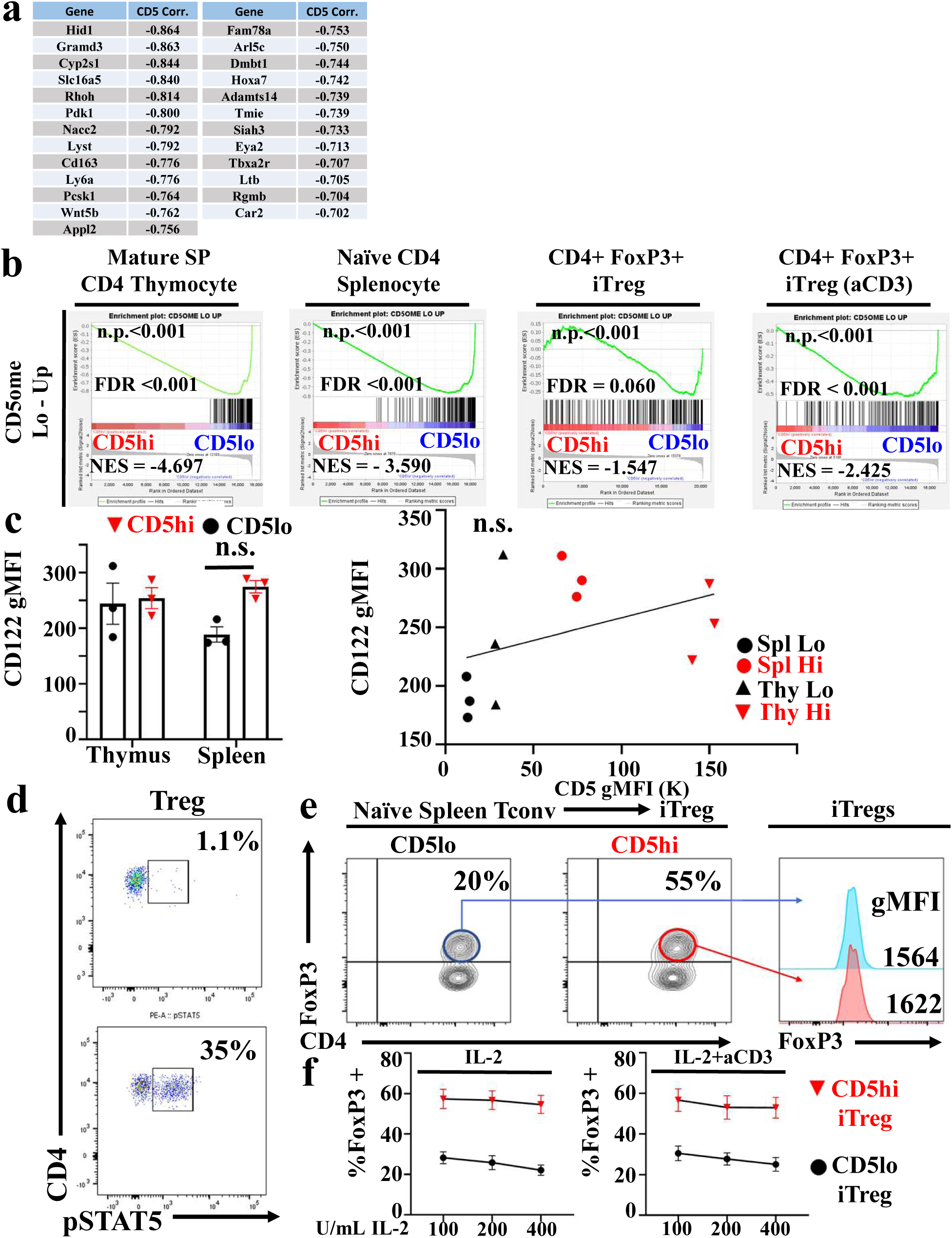
a. Top most CD5 strongly negatively correlated genes across all RNA expression samples b. GSEA plot of 200 most negatively correlated to CD5 genes in gene expression comparisons between CD5hi and CD5lo cells in CD4 single positive (SP) thymocytes, naïve splenocytes, iTregs and iTreg stimulated with aCD3 (left to right). c. CD122 (IL-2Rb) expression on naïve T cells from thymus and spleen (n=3). Data representative of multiple experiments (left). Correlation between CD5 and CD122 expression (right). d. Representative phospho-STAT5 staining on nTregs: Control (top), 30 minutes of IL-2 (bottom). e. Representative comparison of FoxP3 gMFI levels in iTregs from CD5lo and CD5hi Tconv (soluble aCD3 [10ug/mL], IL-2 [50U/mL], and TGFb [2.5ng/mL]). Foxp3 expression in sorted iTregs (OTII/RAGKO/FOXP3-hCD2), 3 days after induction, cultured with variable amounts of IL-2 with or without soluble aCD3.

